# Zebrafish embryos are robust to mechanical perturbations

**DOI:** 10.64898/2026.04.15.718700

**Authors:** Ming Hong Lui, Alejandro Jurado, Leon Lettermann, Timo Betz, Bart E. Vos

## Abstract

The investigation of morphogenesis during organism development has massively benefited from the interaction between developmental biology and biophysics, which gave new quantitative insights into nature’s physical working principles. The process of zebrafish epiboly provides a beautiful model system as cells exhibit collective migration, meanwhile requiring symmetry breaking for gastrulation and the subsequent tissue differentiation. These elegantly robust processes are facilitated by both biochemical and mechanical cues.

Here we test the hypothesis that the future body axis is robust against mechanical perturbations of tissue flow and tissue stress fields. For this, we trigger tissue flow by photoablation, which leads to an externally enforced symmetry breaking before natural symmetry breaking of gastrulation occurs. Using 3D light sheet microscopy, we obtained nuclei trajectories as a proxy for the migration of individual cells. Analyzing the derived tissue velocity field, we observed that epiboly is highly robust and mostly unaffected by ablation damage. However, the photoablation induced a convergent motion of cells towards an azimuthal angle resembling shield formation. We extensively characterized this convergent motion and checked if this tissue flow is able to reorient natural symmetry breaking during shield formation. While ablations can transiently disrupt or reorient the convergent motion, we cannot confirm any redirection of the location where the future body axis of the animal develops. The observations were also highly dependent on the developmental stage of the embryo, which cast a greater and more permanent impact, suggesting strong control exerted by factors other than mechanical signals.

**Significance statement:** Morphogenesis emerges from the interplay between biochemical patterning and mechanical forces, yet the extent to which mechanical cues influence embryo developmental remains unclear. Using zebrafish epiboly as model system, we imposed controlled mechanical perturbations through targeted photoablation and tracking resulting 3D cell movements. We show that although ablations generate transient convergent motions, epiboly is remarkably robust to externally induced tissue flows. Our findings indicate that the future body axis is strongly buffered against mechanical disturbances, highlighting the dominant role of developmental stage and non-mechanical cues in guiding development.

## Introduction

Symmetry-breaking events are essential in the development of an embryo, generally preceding spatial organization such as polarization guiding body axes^1^ and cell differentiation. Here one of the most prominent events is gastrulation, a key process during embryogenesis known to trigger the germ layer differentiation.^2–4^ From a conceptual point of view, characteristic and well-orchestrated changes in collective cell motion are directly coupled to fundamental biological changes of the level of individual cells, a process well described as emergent behavior.^5^ The study of such emergent behavior promises to better understand how complex and highly dynamic natural processes can simultaneously remain robust and well controlled.^6^

The zebrafish is an often-used model organism for studying embryonic development. At 4.3 hours post fertilization (hpf), the blastoderm of a zebrafish embryo contains approximately 2000 cells where the so-called deep cells remain in the interior of the tissue, while a well-defined outer enveloping layer (EVL) of cells forms the interface to the environment. At this time point, epiboly starts, which is the first coordinated movement of cells from the blastoderm towards the yolk^7–9^ (Figure A). This key process is measured in the percentage of cells covering the yolk; for example 50% epiboly refers to the blastoderm cells covering the upper hemisphere of the spherical embryo. Epiboly ends with the full enclosure of the yolk by the developing organism.

At around 5.3 hpf, corresponding to 50% epiboly, two symmetry-breaking events arise. These lay the groundwork for cell differentiation and morphological changes.^7^ First, invagination is observed, where parts of the deep cells underneath the EVL move against the direction of epiboly and invaginate inward and upward. This folding event leads to germ layers with different cell fates and functions.^10, 11^ Second, a shield of thickened cells forms on one side of the sphere, breaking the rotational symmetry, which amplifies the morphological change over time, culminating as the spine of the fish and defining the ventral and dorsal sides^7^ (Figure A).

Over the years, biochemical signals necessary for many morphological changes have been identified.^1, 12–15^ In the context of zebrafish development, epiboly requires coordinated efforts between cytoskeleton, adhesion proteins, but also kinases and transcription regulators.^9^ Additionally, researchers begin to appreciate the interplay between these biochemical organizers and their downstream mechanical consequences such as force transmission, fluidity and surface tension.^16–18^

From a physics point of view, collective cell motion during embryonic development has been modeled using physical frameworks such as percolation theory and fluid dynamics.^19–22^ While these physical models provide general explanations for epiboly motion, cells were assumed to be functionally homogeneous and thus make these models insufficient to elucidate symmetry-breaking events, which could arise from local forces. Consistent with this view, it was shown that differential cortical tension between neighboring cells is sufficient to drive robust cell sorting during zebrafish gastrulation, demonstrating that local variations in mechanical properties can organize tissues even in the absence of explicit biochemical patterning.^23^ In addition, cell division was shown to align with anisotropic tension.^24^

One effective way to study the contribution of local cells and their motion to the overall development of an embryo is to locally induce an external perturbation, such as removal of local tissue. This method dates back to the pioneering studies by Spemann and Mangold, where dorsal cells were transplanted from one newt (*Triton taeniatus*) embryo to another, resulting in gastrulation initiating from the transplanted site.^25^ Later microsurgery studies demonstrated that neural induction is already well underway by the early shield stage.^26^ Modern disruption techniques usually involve photomanipulation: the use of a targeted laser to ablate a precise section of the embryo, and to measure the response afterwards. This can be performed on a small, subcellular scale, for example tracking the tension recoil on the actomyosin ring at the blastoderm-yolk-interface of zebrafish,^11^ or on a tissue scale to study the coordinated responses of the entire blastoderm to greater damage, such as reorientation of the development centre on amniote embryos.^20, 27^ Additionally, photoactivation of cytotoxic proteins is another tool to locally disrupt embryonic development.^28, 29^ However, the precise interplay between local mechanical disruptions and the robust nature of epiboly is still not fully understood.

In this paper, we hypothesize that on an organism level, embryo development should be robust enough to mitigate externally-induced damage at early stages. As a combination of biochemical and mechanical cues is believed to be required for symmetry breaking events, mechanical disruptions alone should not cause permanent disruptions to the developmental trajectory of embryos.

To study global adaptations of the developing embryo, we introduced mechanical perturbations by removing a small section of the enveloping layer of the developing embryo, aiming to elicit mechanical movements of adjacent cells towards the ablation site, akin to shield formation (Figure B). By tracking individual cells with light sheet microscopy, we could quantify collective motion in terms of a coarse-grained velocity field, allowing us to identify the robustness of embryo development under mechanical perturbations.

**Figure 1:**
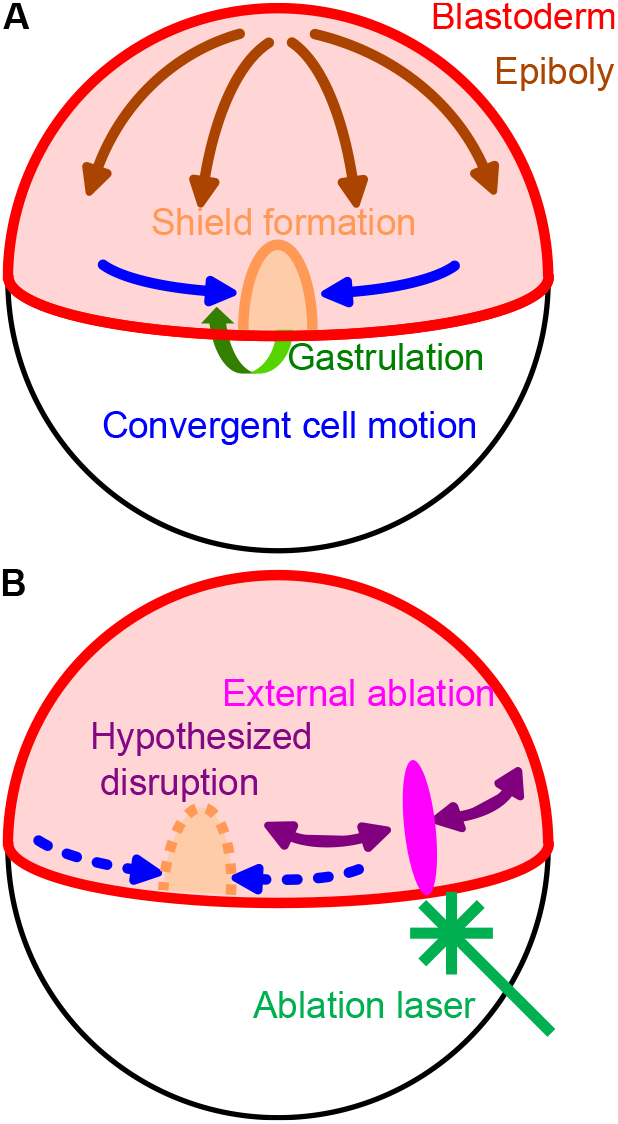
**A**: Normal development of the embryo, where at 50% epiboly invagination sets in and the convergent motion of cells towards the shield region leads to an accumulation of material. **B**: Schematic representation of the hypothesized response of the developing embryo to mechanical perturbations. We hypothesized that with external ablation, it may disrupt flows of the locally adjacent cells, but the overall epiboly or shield formation should not be altered.

## Results

24 transgenic zebrafish embryos (Tg(*β*-actin:H2AmCherry)) with fluorescent histone subunit 2A, representing the cell nucleus as a proxy for cell position,^30^ were imaged in a multiview light sheet microscope equipped with an ablation laser (Figure 2A-B), starting at 4 to 6 hpf until 24 hpf with a temporal resolution of 180 s. To explore the effect of mechanical disruption, we performed laser ablation on 24 embryos 30 minutes after the acquisition began. We also imaged 6 control embryos without ablations to capture the endogenous dynamics of epiboly and symmetry breaking events during development. The consequent tissue flow can be extracted from recording the cell trajectories after the ablation event.

**Figure 2:**
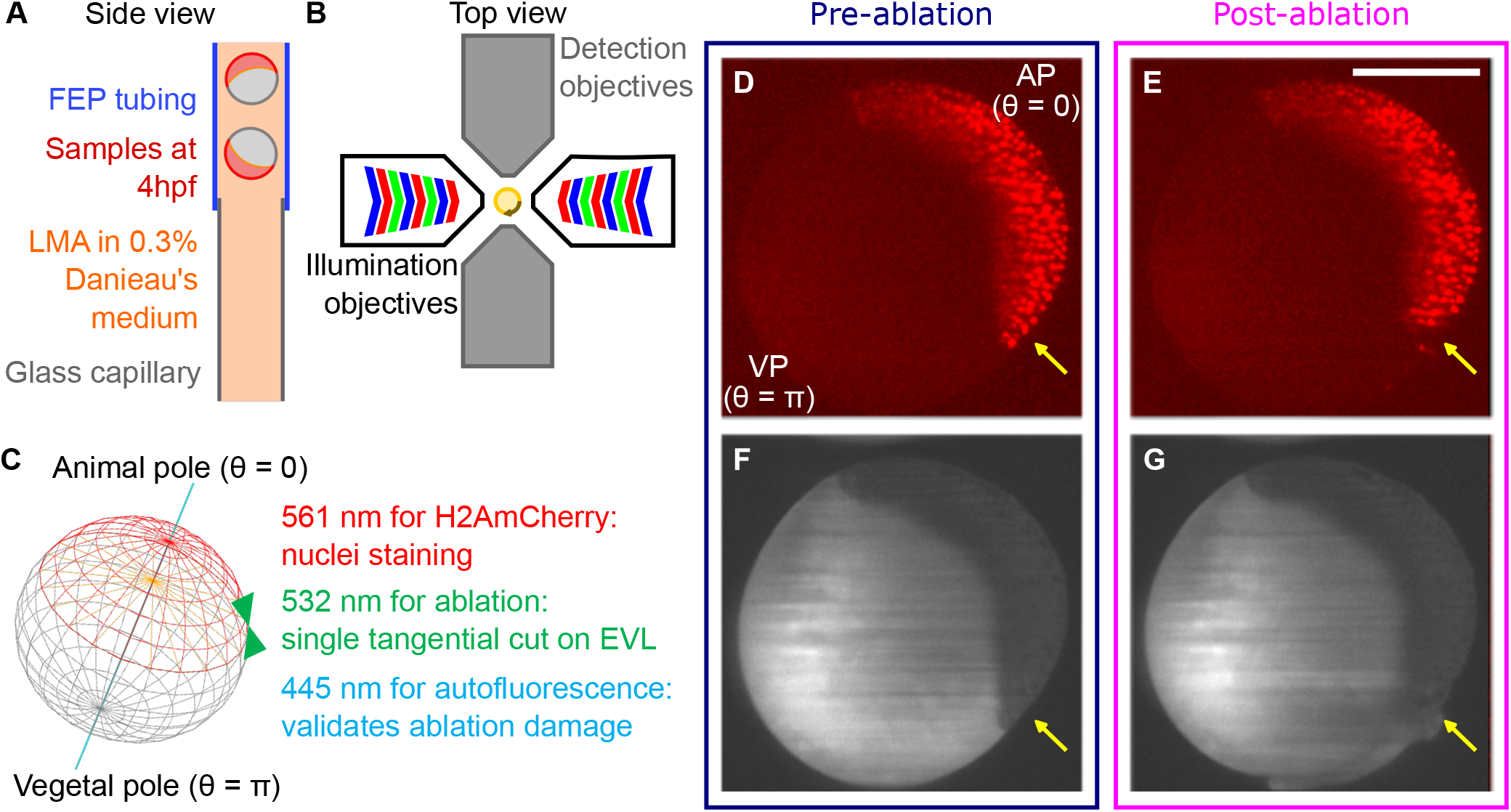
Schematic representation of the experiment. **A**: Zebrafish embryos were loaded and trapped in a transparent cylindrical sample holder. **B**: Setup of a multi-view light sheet microscope. The sample holder was rotated 90 degrees once per time point to provide four acquisition angles for fusion. **C**: Laser ablation was performed tangential to the enveloping layer (EVL), as indicated by the green arrows. Fluorescence signal of a typical experiment of the nuclei before **D** and after **E** ablation is shown, indicating the effect of the ablation. **F-G** show the same embryo in autofluorescence. Scale bar: 200 *µ*m.

Ablations were aligned with the orientation of the embryo, along the direction of epiboly and tangentially across the EVL (Figure 2C). Figure 2D-G show a successful ablation, where only cells local to the ablation site were removed. In addition to the fluorescence signal of the nuclei, the autofluorescence signal of the yolk was recorded to confirm the ablation (Figure 2F-G). All ablations were non-lethal, and embryo development proceeded normally up to 24 hpf without major defects.

### Velocity field captured symmetry breaking events

To quantify the effect of ablation on embryo development, nuclei trajectories were reconstructed from the raw image volumes using TGMM (Tracking with Gaussian Mixture Model),^31^ generating a spatiotemporal lineage of cells for each embryo to infer the velocities of each cell. In some trajectories, we observed that ablations could cause transient and localized disruptions of epiboly of varying magnitudes. Shortly after ablation, adjacent cells moved locally towards the ablation site, potentially for wound healing, while on longer time scales, epiboly proceeded as normal (Figure 3A). However, in other embryos the ablation had little to no effect on the local cell trajectories.

**Figure 3:**
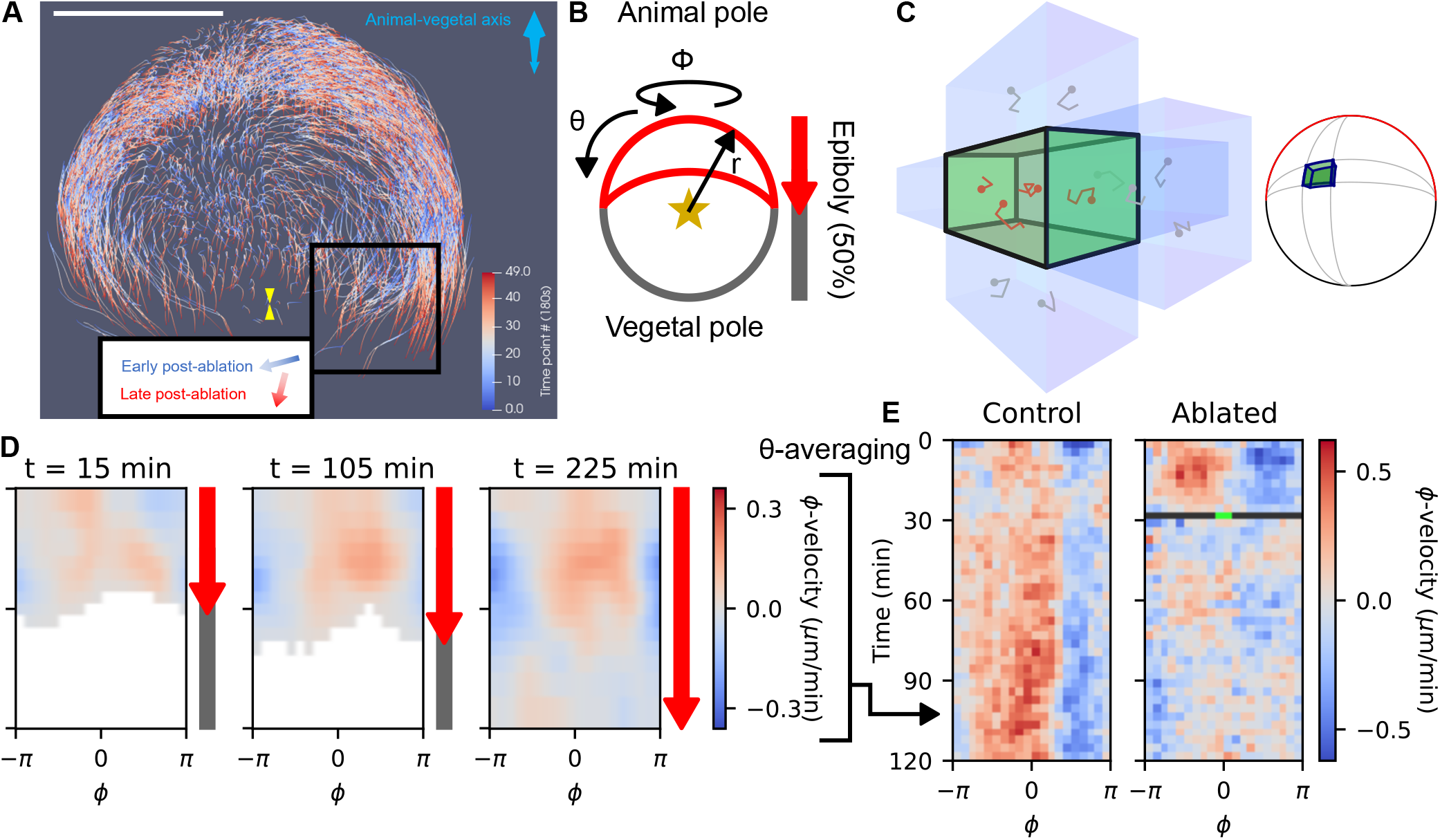
Embryo trajectories and the inferred velocity fields. **A**: TGMM-inferred trajectories of the same embryo after the ablation. Within the first hour (20 time points) after ablation (blue-white gradient), motion of ablation-adjacent cells responded with motion towards the ablation site. Afterwards (white-red gradient), motion was generally towards the vegetal pole as epiboly proceeded as normal. A yellow marker indicates the position of ablation. Scale bar: 200 *µ*m. **B**: Spherical coordinates used in the spherical projection of the embryo. Under this coordinate system epiboly progresses with increasing *θ*. **C**: Viewguide of the construction of the velocity field. Adjacent nuclei trajectories in spatial proximity were binned together and the average velocity was calculated. **D**: Snapshots of the *ϕ*–velocity at different time points, capturing the shield formation as reflected by the convergent motion along the equatorial *ϕ*-bins. **E**: The left shows the same velocity field as **(D)**, now further averaged along the *θ*-direction and introducing the temporal component in the vertical direction. For comparison, the velocity field for an ablated sample is shown on the right. The black line is the time point of ablation, with the green spot the ablation site.

To quantify these observations and facilitate comparisons across samples, we used a spherical projection of the cells’ position that is characterized by a polar and azimuthal angle *θ* and *ϕ* respectively (Figure 3B), such that an increasing angle *θ* points in the direction of epiboly. The vegetal pole (VP) is located at *θ* = *π*, while the animal pole is located at *θ* = 0. Furthermore, the radius *r* indicates the distance to the center of the egg. To reduce the position noise of individual cell movements, we further discretized the approximated sphere into smaller volumetric bins (see Methods), such that velocities of the cell were locally averaged to reconstruct a coarse-grained global velocity field (Figure 3C), similar as done recently.^32^

Figure 3D shows the *ϕ*-velocity for three different timepoints as a function of *θ* and *ϕ*. The systematic emergence of such lateral velocity components is reminiscent of an azimuthal symmetry breaking. Especially the change between the red (positive) to the blue (negative) flow velocities marks a convergence zone towards a defined azimuthal shield angle *ϕ*_0_. This is known to become a region of high cell density and increased thickness (shield formation) that represents the future body axis of the embryo.^33^ Until this point the spherical cap was symmetric with respect to the azimuthal angle *ϕ*. While at early timepoints (left panel) the final convergence zone is not yet clearly defined, later during development a clear preferential angle *ϕ*_0_ emerged, coinciding with shield formation occurring typically at 50% epiboly at 6 hpf^7^ (Supplementary Movie 1). Another symmetry breaking pattern, invagination, could also be captured by the velocity field. During invagination a drastic reorientation of some cells’ motion is observed. Part of the cells that initially move towards the vegetal pole ingress deeper into the embryo towards the yolk, and change direction by moving towards the animal pole. This drastic change in *θ*–velocity is best seen when plotting this velocity component along *r* instead of *ϕ* (Supplementary Movie 2).

To further characterize the temporal dynamics of shield formation, an entire experiment is summarized in Figure 3E by averaging single time points over all *θ*-values. This reduction of dimensionality is justified by a stable position of the convergence zone at *ϕ*_0_ along *θ*. We observed for a control embryo that the position of *ϕ*_0_ is stable and becomes more pronounced in time, while ablation temporarily disrupts the convergent motion.

### Spatiotemporal fitting of global azimuthal velocity to describe shield formation

While analyzing the effect of ablation, the azimuthal convergence zone provides a prominent parameter to further register and align the embryos. Hence, to quantify asymmetric cell migration during embryo development further, we phenomenologically fit a sinusoidal function *A* sin[*ϕ* − *ϕ*_0_(*t*)] to each time point of the averaged azimuthal velocity in Figure 3E (see Figure 4A for an example of the fit), with a fitted phase angle *ϕ*_0_ corresponding to the center of the convergence zone. This is taken to represent the future shield area. A good sinusoidal fit matched closely the convergent motion along the *ϕ*-direction. In contrast, disruptions in the shield formation would lead to a poor sinusoidal fit and unstable phase angles. Tracking the changes in *ϕ*_0_ over time provided a description for the evolution of shield formation.

**Figure 4:**
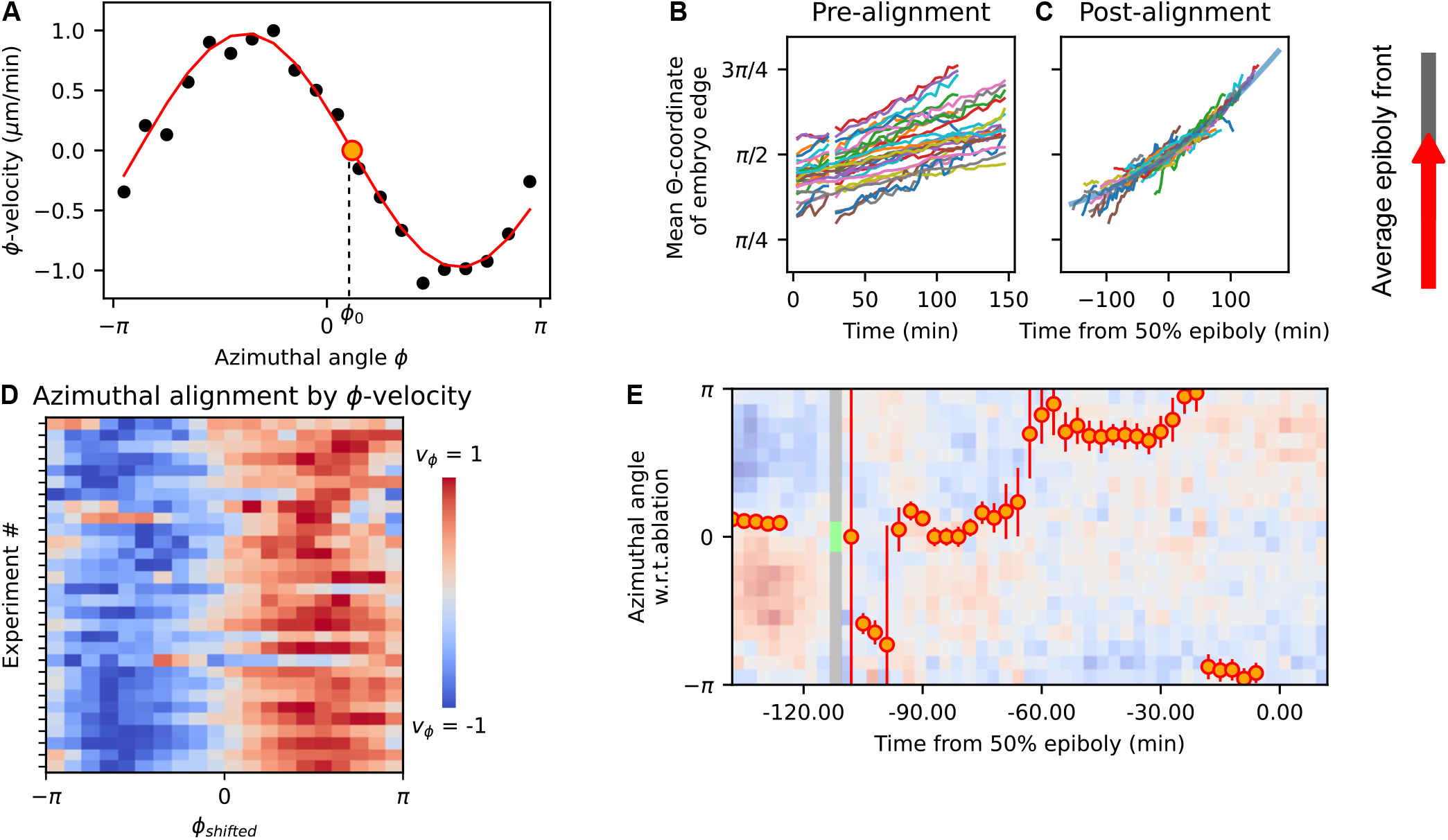
Spatiotemporal alignments of ablation and development. **A**: An example of a negative sinusoidal fit for the *ϕ* velocity to locate the azimuthal angle of convergence *ϕ*_0_. **B-C**: Temporal alignment of all embryos according to epiboly progress. Each experiment was assigned a time offset relative to 50% epiboly. A quadratic curve (thick blue line) was used to align individual measurements in **(C).** The break is caused by the ablation event. **D**: All time-averaged trajectories projected onto the equator for all samples, aligned according to *ϕ*_0_ and with the amplitudes individually normalized. **E**: An example of a *ϕ*–velocity field of an individual embryo aligned spatially and temporally. The velocity field was overlaid with the centres of the sine fit at each time point. Error bars indicate the standard deviation (the square root of the diagonal of the covariance matrix) of the 5-time-point averaged fit. Two quantities, the convergence angle shift relative to the final orientation of the embryo and the standard deviation of the sine fit, were calculated for each time point and compared across samples in Figure 5.

Finally, to compare different embryos, we registered them along the temporal axis. When comparing between experiments, we observed minor variations in the developmental stage prior to imaging among embryos, evidenced by the varying progress of epiboly. Here, based on the coarse-grained velocity field, epiboly is quantitatively defined as the (time-dependent) maximum angle with non-empty bins, averaged across the *ϕ*– and *r*-direction (Figure 4B). To find a temporal alignment for developmental progress based on the phenomenology of epiboly, we assumed a simple quadratic time dependence for epiboly and used the Python-library JAX Autograd to find the best-fitting coefficients using pooled experiments.^34^ Translating the time axis relative to the moment corresponding to 50% epiboly, we could project all measurements onto a single master curve (Figure 4C), showing a good phenomenological fit despite known deviations from simple quadratic behavior, such as the transient slowdown of epiboly at the onset of gastrulation.^9^

Applying sinusoidal fits on the time-averaged *ϕ*-velocities after ablation instead of single time points resulted in good fit qualities for all the experiments. This confirms not only the overall approach, but also that the convergence angle *ϕ*_0_ for shield formation is a robust and universal process in development, regardless of the transient disruption by ablation (Figure 4D). This is further backed by the observation that despite the ablative injury, all but one fish continued to develop for at least 24 hpf.

### Developmental time is a confounding factor for disruptions caused by ablation

Combining all the alignment strategies, we can now describe the azimuthal velocity fields from individual measurements within a common reference frame (Figure 4E), from which we investigated the disruptive effects of our ablations using two derived metrics. First, we measured potential realignment of the convergence angle *ϕ*_0_ after ablation. A change of *ϕ*_0_ after ablation indicates induced instabilities over time and thus long-term changes in the developmental program. Second, to assess the disruption of the convergent motion, we obtained standard deviation (the square root of the diagonal of the covariance matrix) as the uncertainty of the fitted phase angle *ϕ*_0_ of the sinusoidal curve. These two metrics are visualized in Figure 5A and B, respectively.

**Figure 5:**
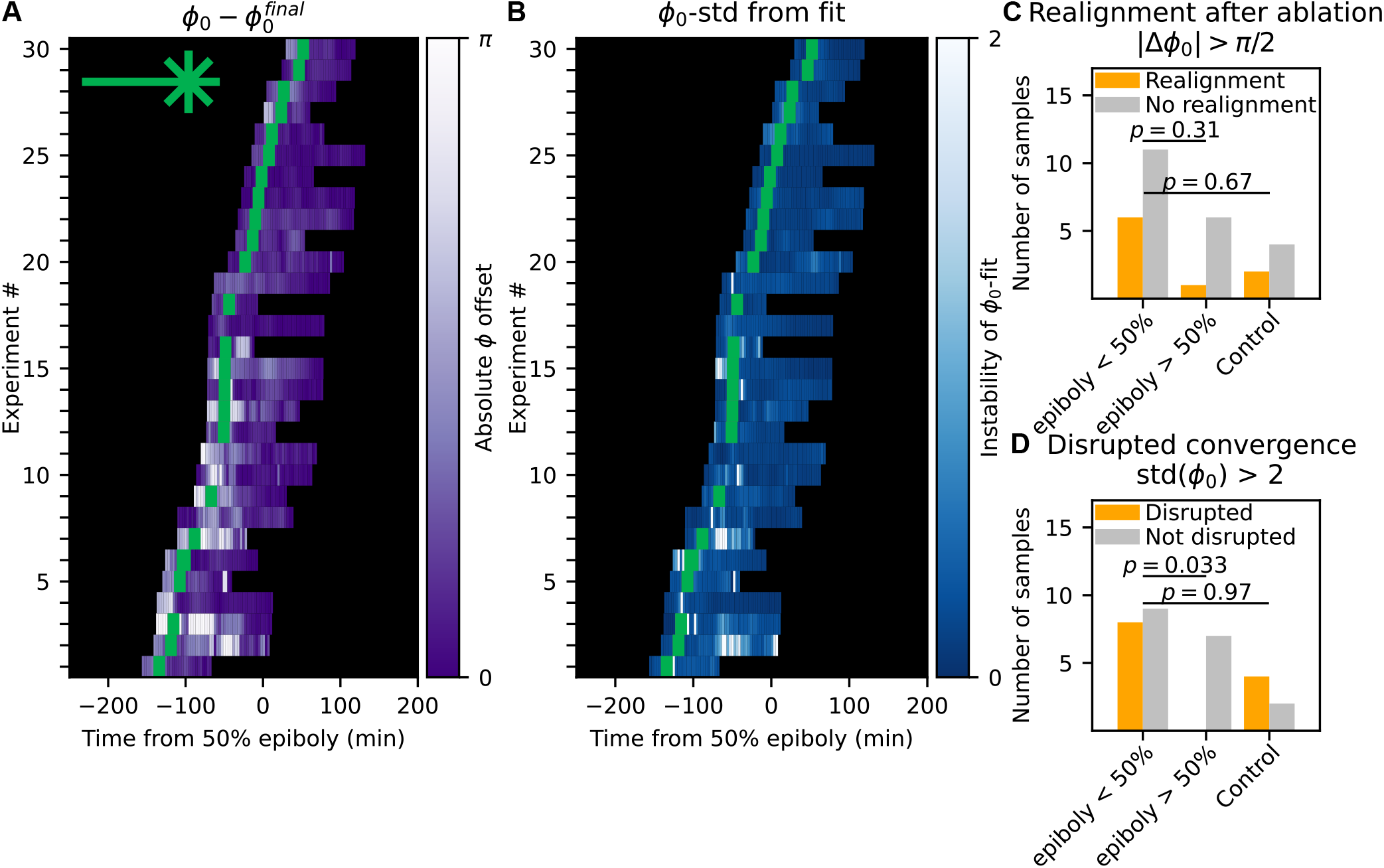
Effect of external disruptions on shield formation confounded by epiboly progress. **A**: Deviation of the angle of convergence for all trajectories, after ordering according to temporal alignment (Figure 4B-C). A brighter color indicates a larger deviation. The green area indicates the ablation event. **B**: standard deviation (the square root of the diagonal of the covariance matrix) of the fitting of *ϕ*_0_ for all trajectories, after ordering according to temporal alignment (Figure 4C-D). **C-D**: Statistics for the occurrences of alignment events, as defined by observing at least one time point with an absolute difference of |Δ*ϕ*_0_| *> π/*2, and statistics for the occurrences of disrupted convergence events, as defined by observing at least one time point with a fitting standard deviation of *>* 2 for *ϕ*_0_. Measurements were grouped by the time of ablation occurring. All *p*-values were calculated using a one-sided Fisher’s exact test.

Figure 5A shows how *ϕ*_0_ changed with respect to its final value, where bright colors indicate large deviations from the final phase angle and dark colors represent stable values. Although there is sample-to-sample variability, sorting the samples by development time relative to 50% epiboly revealed a qualitative trend, with strong changes in *ϕ*_0_ mostly occurring during early development (before 50% epiboly). At later development stages, after 50% epiboly, *ϕ*_0_ has been relatively stable.

As mechanical forces can trigger or disturb the location of gastrulation in *Drosophila*,^35–37^ we wondered if the flow induced by the wound healing after the cut could change the position of invagination in zebrafish. Since we can only retrospectively reconstruct the position of shield formation, the cut is inherently done at a random position. If the convergence flows induced by the wound healing would move the position of the future shield formation, we would expect that the convergence zone that develops after the ablation is close to the final convergence zone, where we define small changes as the difference between the average *ϕ*_0_ for the last 5 and the first 5 time points 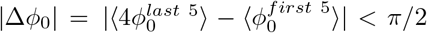. As 50% epiboly marks the onset of invagination and shield formation,^7^ we tested the effect of ablation before and after this time point. If we find a statistical difference with an increase in realignment for ablations done before 50% epiboly, this would suggest that the wound-induced flows have a significant effect on the location of the shield formation. We performed a one-sided Fisher’s exact test between the two groups (Figure 5C) but did not find a statistically significant difference (*p* = 0.31) whether the cut was done before or after shield formation had already been started. This indicates that we could not detect that the flow created by the ablation wound closure disrupts the shield formation, making the position of the body axis highly stable.

In addition to the phase angle *ϕ*_0_, we also obtain the standard deviation of the *ϕ*_0_-fit (Figure 5B). A small standard deviation suggests strong and coherent migration of cells towards *ϕ*_0_ azimuthally, while larger values indicate that there is still uncertainty about the preferential direction of cell migration and weaker organization. Again, sorting by time revealed much stronger uncertainty before 50% epiboly than after.

Comparing the two groups of ablated samples before and after 50% epiboly, we observed significant differences (Fisher’s exact test *p* = 0.033) in the proportion of samples with transient uncertainty, defined by at least one time point during acquisition with a standard deviation *std*(*ϕ*_0_) *>* 2 (Figure 5D). This suggested developmental time is likely a confounding factor for the coherence of collective motion.

Regarding the effect of laser ablation, when comparing between the samples ablated before 50% epiboly and the control samples, all of which started acquisition before 50% epiboly, we did not observe significant differences both in terms of the proportion of samples experiencing realigned or disrupted collective motion (Figure 5C and D). This suggested that external ablation did not have a significant effect under the confounding influence of developmental time. The embryos seemingly have robust programs regarding orientation of shield formation, and were robust to external, localized disruptions. This is consistent with the current picture of morphogen-driven gastrulation and shield formation.

## Discussion

### On the robustness of shield formation

In most embryos studied here, we observed a transient loss of azimuthal directionality immediately after laser ablation that was reminiscent of cells flowing in the ablated region. However, already within 30 minutes, the convergent flow was restored back to normal epiboly motion (Figure 5D). Although we cannot rule out a tensional force driving the tissue flow, we did not observe an immediate recoil indicative of pre-existing tensile stress at the ablation site. However, it remains possible that the time resolution of 3 minutes provided by the 3D light sheet microscope is not sufficient to detect such an effect, as the wound healing might already start within seconds after wounding. Still, the observed changes in flow upon ablation could be attributed to the disrupted surface tension of the EVL which is known to be maintained by actomyosin flows at the yolk syncytial layer during epiboly^38^ and propagated via cell-cell adhesion.^39^ Ablative damage is expected to drive a transient redistribution of mechanical stress before the tissue integrity was eventually restored.

Although we observed a change in the azimuthal angle of convergence *ϕ*_0_ in some embryos after the ablation, we did not see a statistically significant difference for such occurrences between ablated embryos and the controls. While limited by sample size, the simplest explanation for this is that mechanical ablation alone is insufficient for altering the trajectories of the cells that eventually results in shield formation. There seems to be strong robustness in the pre-programmed developmental trajectory, as most realignment occurred before 50% epiboly, again irrespective of ablations (Figure 5B).

Such robustness may come from pre-established biochemical organizers,^8, 40–43^ and the interplay between mechanical movement and chemical signals should be emphasized. We have also noted that goosecoid, expressed in cells at the dorsal midline,^33^ aligned with the angle of azimuthal convergence (Figure S4). More recently, migrasomes, a novel organelle forming at the shield, have been discovered to provide potential morphogenesis cues.^44^ The expression of these biological organizers begins prior to 50% epiboly, after which expression plateaus. This aligns well with our observed uncertainty of azimuthal convergence before 50% epiboly (Figure 5B,E). These mediated signals, unaffected by EVL damage, could help maintain the robustness of shield formation in our experiments, suggesting that the dorsal-ventral symmetry breaking is dominated by biochemical rather than mechanical signals, and azimuthal convergence of cells is an effect, rather than the cause, of shield formation.

However, we must acknowledge the possibility that stronger, lethal ablations could have a more persistent effect for realignment, and a survivor bias here meant we are limited to studying the effects of localized tension disruption, which apparently could not steer shield formation. Nonetheless, if realignment via mechanical forces is unattainable without lethality, the above argument of biochemical signals driving shield formation still stands as shield formation is a naturally occurring event in embryos.

### On the time-dependent progression of epiboly

In this work, we have demonstrated that velocity fields provide a framework to quantify symmetry-breaking in early embryonic developmental stages, and used various downstream analyses to characterize gastrulation and shield formation in zebrafish. Recent studies have also used velocity fields to analyze global coordinated wave motions in cellular spheroids to infer velocity field patterns such as vortices and global rotations.^19, 45^ This quantitatively connects experimental observations with theoretical predictions to study underlying forces driving symmetry breaking events.

A widely cited feature in zebrafish development since its first report by Kimmel et al.^7^ is the so-called epibolic pause. At 50% epiboly, EVL movement is reported to halt for up to one hour at the onset of ingression. Kimmel et al. based their report on an idealized observation performed on a stereomicroscope, which later became a well-established reference in literature.^9, 46, 47^

However, the tracks of fluorescently labeled nuclei in our study reveal that a small group of cells at the leading edge advanced uninterrupted throughout epiboly with a constant velocity until ≈ 50% epiboly, and even accelerate afterwards (see Figure 4B). This, to our best knowledge, has not been described in literature, and the cause for this discrepancy is likely found in the high resolution capabilities of light-sheet microscopy combined with fluorescence. These data suggest that the apparent pause reflects the bulk of the tissue, rather than a complete halt of movement. From a mechanical perspective, an acceleration after 50% epiboly is consistent with actomyosin-driven tension during gastrulation.^38^ Our findings therefore call for a refined, scale-dependent interpretation of epiboly dynamics.

## Conclusion

In this study we investigated the robustness of the developing zebrafish embryo against mechanical perturbations induced by laser ablation. By tracing the individual nuclei, we observed the spreading of cells over the surface of the yolk. A mechanical perturbation, induced by laser ablation, could cause a temporary loss of organization of cellular movement, but eventually migration recovers its directionality, consistent with the known robustness of developing zebrafish. There is an apparent correlation between the azimuthal velocity convergence and developmental time, suggesting the versatile pre-programming of the shield formation process.

## Materials and Methods

### Zebrafish embryo imaging

Transgenic female zebrafish Tg(*β*-actin:H2AmCherry) were crossed with wildtype AB males. Fertilized eggs were grown in 0.3% w/v Danieau’s medium (19 mM NaCl, 0.2 mM KCl, 0.1 mM MgSO4, 0.2mM Ca(NO3)2, 1.7mM HEPES) at 28°C and mechanically dechorionated at 2 hpf. At 4 hpf, embryos attained a spherical geometry. The embryos were trapped two at a time in a segment of fluorinated ethylene propylene (FEP) tubing with a 2 mm outer diameter appended to a glass capillary (Blaubrand 708744), using 1% low melting agarose (LMA) dissolved in the same medium. The whole sample was rotated for several minutes along the capillary axis to ensure embryos remained fixed in a central position, avoiding them touching the FEP walls. Once LMA was fully gelified, samples were stored fully submerged in Danieau’s water at 28°C to avoid drying.

### Lightsheet Microscopy with ablation

To capture the entirety of the embryonic volume, we deployed multiview selective plane illumination microscopy (SPIM), using Luxendo MuVi SPIM (Bruker Corporation) for acquisition, image registration and orthogonal fusion for the final volume. We used an excitation wavelength of 561 nm to image the H2AmCherry signals corresponding to nuclei fluorescence, and an excitation wavelength of 445 nm to capture autofluorescence of the embryo to confirm the ablation damage. After sample mounting, the center of each embryo was registered as the pivot for sample rotation and four orthogonal views of the whole volume were acquired every 180 seconds. Views were acquired two at a time using opposing cameras (see Figure 2), meaning a single rotation was enough to cover all angles. Luxendo post-processing software was used to register all views in a single .hd5 file, rendering isotropic resolution with voxels of 1 *µm*^3^. The microscope was equiped with an additional 60 mW pulsed, high energy, 532 nm ablation laser, used for simultaneous ablation while imaging the embryo. The ablation was tangential to the enveloping layer, in the direction of epiboly, consisting of 41 pulses spaced 1.2 µm apart. For the whole duration of the experiment, embryos were imaged inside an incubation chamber fully submerged in Danieau’s water at 28°C, maintaining physiological conditions.

### Velocity field construction from imaging data

Nuclei trajectories were extracted from the mCherry image stack using TGMM.^31^ The trajectories were calculated separately before and after the ablation. To reduce noise from the individual trajectories, the embryo geometry was coarse-grained into volumetric bins following a spherical approximation. In brief, the radius and centre of the sphere was determined by least square fitting all pre-ablation trajectories onto a spherical shell irrespective of time, and embryos were rotated to have the blastoderm center of mass pointing upwards in spherical coordinates, such that the animal pole is located at *θ* = 0 and the vegetal pole at *θ* = *π*. A further rotation along this newly defined *θ* axis was performed to align all ablation sites to *ϕ* = 0. After these rotations, epiboly progression can be easily quantified as the leading edge of cells along the *θ* angle, with 50% epiboly at 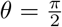. Furthermore, the ablation site is aligned for all embryos.

Exact temporal referencing of different experiments is practically difficult since Zebrafish females can release several batches of eggs consecutively which are later indistinguishable even with temporal registration. For this reason, a minimal temporal registration was needed (see Figure 4B) to correct for temporal shifts. The leading edge of epiboly was taken as a proxy for developmental time and *θ*_*lead*_ − *time* curves were time shifted to collapse onto an epiboly master curve.

In the new coordinate system, the sphere was subdivided into smaller volumetric bins in *ϕ, θ* and *r*. In total, considering the density of occupation of cells, the chosen subdivisions were respectively *n*_*ϕ*_ = 20, *n*_*θ*_ = 15 and *n*_*r*_ = 11, resulting in a mesh-grid of 3300 bins. Notably, radial subdivision spanned between −80 *µm < r <* 65 *µm* relative to the average radial distances of all nuclei, and the polar division was proportional to an inverse cosine function such that bins near the poles have compensatory larger volumes.

Velocities of the nuclei were inferred using a four-point average (or two-point if at both ends of the trajectory). Individual velocities were averaged within each bin. Dimensionality was further reduced by averaging the velocity across multiple bins along spherical coordinates or along time, or by decomposing the average velocity in *r*-, *θ*- or *ϕ*-components.

### Statistical analysis and data representation

## Data availability

All experimental data is available from the authors upon request.

## Code availability

All code is available from the authors upon request.

## Author contributions

ML collected data. ML and BV analyzed data. AJ and LL wrote the analysis program. TB conceived the study. ML, AJ, TB and BV wrote the paper. All authors reviewed and approved the final version of the paper.

## Acknowledgements

TB and BV were supported by the European Research Council (ERC grant 771201). We thank the Göttingen Center for Molecular Biosciences (GZBM) for the zebrafish facility and animal care. The instrument used was funded by the Deutsche Forschungsgemeinschaft (DFG, German Research Foundation) – Projektnummer 455153711 and 456112451. This research was conducted within the Max Planck School Matter to Life, supported by the Dieter Schwarz Foundation and the German Federal Ministry of Research, Technology and Space (BMFTR) in collaboration with the Max Planck Society.

## Supplementary Material

**Supplementary Movie 1**: the time development of the *θ*-, *ϕ*-, and *r*-velocities as a function of the polar and azimuthal angle *θ* and *ϕ*.

**Supplementary Movie 2**: the time development of the *θ*-, *ϕ*-, and *r*-velocities as a function of the polar angle *θ* and distance to the embryo center *r*.

## References

[1] C. P. Petersen and P. W. Reddien, “Wnt signaling and the polarity of the primary body axis,” Cell, vol. 139, no. 6, pp. 1056–1068, 2009.

[2] V. Hamburger and H. L. Hamilton, “A series of normal stages in the development of the chick embryo,” Developmental dynamics, vol. 195, no. 4, pp. 231–272, 1992.

[3] K. K. Hisaoka and H. I. Battle, “The normal developmental stages of the zebrafish, brachydanio rerio (hamilton-buchanan),” Journal of Morphology, vol. 102, no. 2, pp. 311–327, 1958.

[4] R. Keller, L. A. Davidson, and D. R. Shook, “How we are shaped: the biomechanics of gastrulation,” Differentiation: ORIGINAL ARTICLE, vol. 71, no. 3, pp. 171–205, 2003.

[5] E. Scarpa and R. Mayor, “Collective cell migration in development,” The Journal of cell biology, vol. 212, no. 2, pp. 143–155, 2016.

[6] X. Trepat and E. Sahai, “Mesoscale physical principles of collective cell organization,” Nature Physics, vol. 14, no. 7, pp. 671–682, 2018.

[7] C. B. Kimmel, W. W. Ballard, S. R. Kimmel, B. Ullmann, and T. F. Schilling, “Stages of embryonic development of the zebrafish,” Developmental dynamics : an official publication of the American Association of Anatomists, vol. 203, no. 3, pp. 253–310, 1995.

[8] L. A. Rohde and C.-P. Heisenberg, “Zebrafish gastrulation: cell movements, signals, and mechanisms,” International review of cytology, vol. 261, pp. 159–192, 2007.

[9] S. E. Lepage and A. E. E. Bruce, “Zebrafish epiboly: mechanics and mechanisms,” The International journal of developmental biology, vol. 54, no. 8-9, pp. 1213–1228, 2010.

[10] R. M. Warga and C. B. Kimmel, “Cell movements during epiboly and gastrulation in zebrafish,” Development, vol. 108, no. 4, pp. 569–580, 1990.

[11] B. Alberts, D. Bray, K. Hopkin, A. D. Johnson, J. Lewis, M. Raff, K. Roberts, and P. Walter, Essential cell biology. Garland Science, 2015.

[12] Alan Mathison Turing, “The chemical basis of morphogenesis,” Philosophical Transactions of the Royal Society of London. Series B, Biological Sciences, vol. 237, no. 641, pp. 37–72, 1952.

[13] R. Sheth, L. Marcon, M. F. Bastida, M. Junco, L. Quintana, R. Dahn, M. Kmita, J. Sharpe, and M. A. Ros, “Hox genes regulate digit patterning by controlling the wavelength of a turing-type mechanism,” Science (New York, N.Y.), vol. 338, no. 6113, pp. 1476–1480, 2012.

[14] S. Li, D. Edgar, R. Fässler, W. Wadsworth, and P. D. Yurchenco, “The role of laminin in embryonic cell polarization and tissue organization,” Developmental cell, vol. 4, no. 5, pp. 613–624, 2003.

[15] S. Nair and T. F. Schilling, “Chemokine signaling controls endodermal migration during zebrafish gastrulation,” Science (New York, N.Y.), vol. 322, no. 5898, pp. 89–92, 2008.

[16] P. Tomakidi, S. Schulz, S. Proksch, W. Weber, and T. Steinberg, “Focal adhesion kinase (fak) perspectives in mechanobiology: implications for cell behaviour,” Cell and tissue research, vol. 357, no. 3, pp. 515–526, 2014.

[17] N. I. Petridou, S. Grigolon, G. Salbreux, E. Hannezo, and C.-P. Heisenberg, “Fluidization-mediated tissue spreading by mitotic cell rounding and non-canonical wnt signalling,” Nature cell biology, vol. 21, no. 2, pp. 169–178, 2019.

[18] A. Hernández-Vega, M. Marsal, P.-A. Pouille, S. Tosi, J. Colombelli, T. Luque, D. Navajas, I. Pagonabarraga, and E. Martín-Blanco, “Polarized cortical tension drives zebrafish epiboly movements,” The EMBO journal, vol. 36, no. 1, pp. 25–41, 2017.

[19] B. Wallmeyer, S. Trinschek, S. Yigit, U. Thiele, and T. Betz, “Collective cell migration in embryogenesis follows the laws of wetting,” Biophysical journal, vol. 114, no. 1, pp. 213–222, 2018.

[20] M. Saadaoui, D. Rocancourt, J. Roussel, F. Corson, and J. Gros, “A tensile ring drives tissue flows to shape the gastrulating amniote embryo,” Science (New York, N.Y.), vol. 367, no. 6476, pp. 453–458, 2020.

[21] N. I. Petridou, B. Corominas-Murtra, C.-P. Heisenberg, and E. Hannezo, “Rigidity percolation uncovers a structural basis for embryonic tissue phase transitions,” Cell, vol. 184, no. 7, pp. 1914–1928.e19, 2021.

[22] A. Mongera, P. Rowghanian, H. J. Gustafson, E. Shelton, D. A. Kealhofer, E. K. Carn, F. Serwane, A. A. Lucio, J. Giammona, and O. Campàs, “A fluid-to-solid jamming transition underlies vertebrate body axis elongation,” Nature, vol. 561, no. 7723, pp. 401–405, 2018.

[23] J.-L. Maître, H. Berthoumieux, S. F. G. Krens, G. Salbreux, F. Jülicher, E. Paluch, and C.-P. Heisenberg, “Adhesion functions in cell sorting by mechanically coupling the cortices of adhering cells,” Science (New York, N.Y.), vol. 338, no. 6104, pp. 253–256, 2012.

[24] P. Campinho, M. Behrndt, J. Ranft, T. Risler, N. Minc, and C.-P. Heisenberg, “Tension-oriented cell divisions limit anisotropic tissue tension in epithelial spreading during zebrafish epiboly,” Nature cell biology, vol. 15, no. 12, pp. 1405–1414, 2013.

[25] H. Spemann and H. Mangold, “über induktion von embryonalanlagen durch implantation artfremder organisatoren,” Archiv für mikroskopische Anatomie und Entwicklungsmechanik, vol. 100, no. 3, pp. 599–638, 1924.

[26] J. Shih and S. E. Fraser, “Characterizing the zebrafish organizer: microsurgical analysis at the early-shield stage,” Development, vol. 122, no. 4, pp. 1313–1322, 1996.

[27] P. Caldarelli, A. Chamolly, A. Villedieu, O. Alegria-Prévot, C. Phan, J. Gros, and F. Corson, “Self-organized tissue mechanics underlie embryonic regulation,” Nature, vol. 633, no. 8031, pp. 887–894, 2024.

[28] S. Curado, D. Y. R. Stainier, and R. M. Anderson, “Nitroreductase-mediated cell/tissue ablation in zebrafish: a spatially and temporally controlled ablation method with applications in developmental and regeneration studies,” Nature protocols, vol. 3, no. 6, pp. 948–954, 2008.

[29] K. Mruk, P. Ciepla, P. A. Piza, M. A. Alnaqib, and J. K. Chen, “Targeted cell ablation in zebrafish using optogenetic transcriptional control,” Development, vol. 147, no. 12, 2020.

[30] J. Huisken and D. Y. R. Stainier, “Even fluorescence excitation by multidirectional selective plane illumination microscopy (mspim),” Optics letters, vol. 32, no. 17, pp. 2608–2610, 2007.

[31] K. McDole, L. Guignard, F. Amat, A. Berger, G. Malandain, L. A. Royer, S. C. Turaga, K. Branson, and P. J. Keller, “In toto imaging and reconstruction of post-implantation mouse development at the single-cell level,” Cell, vol. 175, no. 3, pp. 859–876.e33, 2018.

[32] L. Lettermann, A. Jurado, T. Betz, F. Wörgötter, and S. Herzog, “Tutorial: a beginner’s guide to building a representative model of dynamical systems using the adjoint method,” Communications Physics, vol. 7, no. 1, 2024.

[33] C. Thisse, B. Thisse, M. E. Halpern, and J. H. Postlethwait, “Goosecoid expression in neurectoderm and mesendoderm is disrupted in zebrafish cyclops gastrulas,” Developmental biology, vol. 164, no. 2, pp. 420–429, 1994.

[34] J. Bradbury, R. Frostig, P. Hawkins, M. J. Johnson, C. Leary, D. Maclaurin, G. Necula, A. Paszke, J. VanderPlas, S. Wanderman-Milne, and Q. Zhang, “Jax: composable transformations of python+ numpy programs,” 2018.

[35] E. Farge, “Mechanical induction of twist in the drosophila foregut/stomodeal primordium,” Current biology : CB, vol. 13, no. 16, pp. 1365–1377, 2003.

[36] N. Desprat, W. Supatto, P.-A. Pouille, E. Beaurepaire, and E. Farge, “Tissue deformation modulates twist expression to determine anterior midgut differentiation in drosophila embryos,” Developmental cell, vol. 15, no. 3, pp. 470–477, 2008.

[37] T. Brunet, A. Bouclet, P. Ahmadi, D. Mitrossilis, B. Driquez, A.-C. Brunet, L. Henry, F. Serman, G. Béalle, C. Ménager, F. Dumas-Bouchiat, D. Givord, C. Yanicostas, D. Le-Roy, N. M. Dempsey, A. Plessis, and E. Farge, “Evolutionary conservation of early mesoderm specification by mechanotransduction in bilateria,” Nature communications, vol. 4, p. 2821, 2013.

[38] M. Behrndt, G. Salbreux, P. Campinho, R. Hauschild, F. Oswald, J. Roensch, S. W. Grill, and C.-P. Heisenberg, “Forces driving epithelial spreading in zebrafish gastrulation,” Science (New York, N.Y.), vol. 338, no. 6104, pp. 257–260, 2012.

[39] B. D. Crawford, C. A. Henry, T. A. Clason, A. L. Becker, and M. B. Hille, “Activity and distribution of paxillin, focal adhesion kinase, and cadherin indicate cooperative roles during zebrafish morphogenesis,” Molecular biology of the cell, vol. 14, no. 8, pp. 3065–3081, 2003.

[40] S. Schneider, H. Steinbeisser, R. M. Warga, and P. Hausen, “Beta-catenin translocation into nuclei demarcates the dorsalizing centers in frog and fish embryos,” Mechanisms of development, vol. 57, no. 2, pp. 191–198, 1996.

[41] S. T. Dougan, R. M. Warga, D. A. Kane, A. F. Schier, and W. S. Talbot, “The role of the zebrafish nodal-related genes squint and cyclops in patterning of mesendoderm,” Development, vol. 130, no. 9, pp. 1837–1851, 2003.

[42] F. L. Marlow, “Setting up for gastrulation in zebrafish,” Current topics in developmental biology, vol. 136, pp. 33–83, 2020.

[43] D. Pinheiro and C.-P. Heisenberg, “Zebrafish gastrulation: Putting fate in motion,” Current topics in developmental biology, vol. 136, pp. 343–375, 2020.

[44] D. Jiang, Z. Jiang, Di Lu, X. Wang, H. Liang, J. Zhang, Y. Meng, Y. Li, D. Wu, Y. Huang, Y. Chen, H. Deng, Q. Wu, J. Xiong, A. Meng, and L. Yu, “Migrasomes provide regional cues for organ morphogenesis during zebrafish gastrulation,” Nature cell biology, vol. 21, no. 8, pp. 966–977, 2019.

[45] T. Brandstätter, D. B. Brückner, Y. L. Han, R. Alert, M. Guo, and C. P. Broedersz, “Curvature induces active velocity waves in rotating spherical tissues,” Nature communications, vol. 14, no. 1, p. 1643, 2023.

[46] S.-J. Lee, “Dynamic regulation of the microtubule and actin cytoskeleton in zebrafish epiboly,” Biochemical and biophysical research communications, vol. 452, no. 1, pp. 1–7, 2014.

[47] A. E. E. Bruce, “Zebrafish epiboly: Spreading thin over the yolk,” Developmental dynamics : an official publication of the American Association of Anatomists, vol. 245, no. 3, pp. 244–258, 2016.

